# Unravelling the impact of aging on the human endothelial lncRNA transcriptome

**DOI:** 10.1101/2022.09.05.506634

**Authors:** Maria-Kyriaki Drekolia, Sweta Talyan, Rebeca Cordellini Emidio, Reinier Abraham Boon, Stefan Guenther, Mario Looso, Gabrijela Dumbović, Sofia-Iris Bibli

## Abstract

The incidence and prevalence of cardiovascular disease is highest among the elderly. There is a need to further understand the mechanisms behind endothelial cell aging in order to achieve vascular rejuvenation and minimize the onset of age-related vascular diseases. Long non-coding RNAs (lncRNAs) have been proposed to regulate numerous processes in the human genome, yet their function in vascular aging and their therapeutic potential remain largely unknown. This is primarily because the majority of studies investigating the impact of aging on lncRNA expression heavily rely on *in vitro* studies based on replicative senescence. Here, using a unique collection of young and aged endothelial cells isolated from native human arteries, we sought to characterize the age-related alterations in lncRNA expression profiles. We were able to detect a total of 4463 lncRNAs expressed in the human endothelium from which ∼17% (798) were altered in advanced age. One of the most affected lncRNAs in aging was the primate-specific, Prostate Cancer Associated Transcript (PCAT) 14. In our follow up analysis, using single molecule RNA FISH, we showed that PCAT14 is relatively abundant, localized almost exclusively in the nucleus of young endothelial cells, and silenced in the aged endothelium. Functionally, our studies proposed that PCAT14 affects multiple endothelial cell functions including endothelial cell migration, sprouting and inflammatory responses. Taken together, our data highlight that endothelial cell aging correlates with altered expression of lncRNAs, which could impair the endothelial regenerative capacity and enhance inflammatory phenotypes.

## Introduction

Aging is defined as a gradual decline in organism function coupled with marked multi-organ changes in redox balance, DNA integrity, telomere length, mitochondrial function as well as non-coding RNA expression (López-Otín et al., 2013). Recent studies propose that endothelial aging and senescence impacts systemic metabolic changes which result in the development of chronic diseases such as diabetes, obesity and atherosclerosis (Campisi and Di d’Adda Fagagna, 2007; Ting et al., 2021). The impact of vascular aging and vascular decline on the overall youthfulness of tissues is receiving growing attention (Le Couteur and Lakatta, 2010), and interestingly induction of angiogenesis in aged mice rescued detrimental effects of aging, proposing that preserving endothelial cell regeneration can serve as a rejuvenation strategy (Das et al., 2018). In the same context, the use of senolytics to target endothelial cell senescence was recently proposed as a novel organismal rejuvenation approach (Suda et al., 2021). With the increasing life expectancy, the number of patients with age-related cardiovascular diseases will rise in the near future, leading to an increased healthcare burden. There is a need for new therapies to treat vascular and endothelial cell aging and minimize the onset of age-related diseases.

Currently there are over 16,000 annotated long-noncoding RNAs (lncRNAs) in mammalian genomes (Derrien et al. 2012; Cabili et al. 2011; Consortium 2012; Abascal et al. 2020), which is proportional to the number of annotated protein-coding genes. Over the past two decades, numerous studies demonstrated that lncRNAs can influence gene expression programs, cellular differentiation and organismal development (Mongelli et al., 2019; Rinn and Chang, 2020; Statello et al., 2021) lncRNAs were shown to have important roles in vascular diseases and aging (Bink et al., 2019), regulating endothelial cell cycle control (Tripathi et al., 2013; He et al., 2015; Boon et al., 2016; Hofmann et al., 2019) and inflammatory profiles (Holdt et al., 2010; Cremer et al., 2019). Identification of lncRNAs regulated by aging and studying their downstream molecular targets would potentially unravel novel mechanisms of aging and open new avenues for vascular rejuvenation. Nevertheless, the role of lncRNAs in endothelial cell aging remains largely unknown, in part because most of the studies linking lncRNAs with age-associated diseases and endothelial senescence have been performed in cultured human endothelial cells undergone replicative senescence following multiple duplication cycles or animal models. To address this in this study we mapped the human native endothelial lncRNA expression profiles from young and aged individuals. Collectively, our analysis shows lncRNAs are highly affected by aging in endothelial cells, wherein they may play a causative role in the endothelial dysfunctional phenotypes. Furthermore, we provide a valuable resource for the community to investigate lncRNA expression dynamics in aging of primary endothelial cells.

## Materials and methods Materials

Cell culture media and phosphate buffer saline were from Gibco (Invitrogen; Darmstadt, Germany). Fibronectin was from Corning (Corning GmBH, Kaiserslautern, Germany), interleukin (IL)-1β was from Proteintech (Manchester, UK).

### Endothelial cell isolation and culture

#### Human endothelial cells from mesenteric arteries

Healthy mesenteric arteries from humans of 20±2.3 and 80±3.4 years of age, were isolated as previously described (Bibli et al., 2021).

#### Human umbilical vein endothelial cells

Cells were isolated and cultured as described (Bibli et al., 2021) and were used (passage up to 3) for the different experiments.

#### THP-1 cells

THP-1 monocytic cells were obtained from the American Type Culture Collection (LGC Standards) and cultured in RPMI-1640 containing 2 mmol/L glutamine, 10 mmol/L HEPES, 1 mmol/L sodium pyruvate, 4.5g/L glucose, 1.5 g/L sodium bicarbonate and 10% FBS.

All cells were negative for mycoplasma contamination. Cultured cells were kept in a humidified incubator at 37°C containing 5% CO_2_.

### RNA sequencing

To extract total RNA from endothelial cells, samples were processed with the RNeasy Mini Kit (Qiagen, Hilden, Germany) and subjected to on-column DNase digestion according to the manufacturer’s protocol. RNA and library preparation integrity were verified with LabChip Gx Touch 24 (Perkin Elmer). 4μg of total RNA was used as input for VAHTS Stranded RNA-seq Library preparation following manufacture’s protocol (Vazyme). Sequencing was performed on NextSeq2000 instrument (Illumina) using P3 flowcell with 1×72bp single end setup. The resulting raw reads were assessed for quality, adapter content and duplication rates with FastQC (http://www.bioinformatics.babraham.ac.uk/projects/fastqc). Trimmomatic version 0.39 was employed to trim reads after a quality drop below a mean of Q20 in a window of 5 nucleotides (Bolger et al., 2014). Only reads between 30 and 150 nucleotides were cleared for further analyses. Trimmed and filtered reads were aligned versus the Ensembl human genome version hg38 (Ensembl release 104) using STAR 2.7.9a with the parameter “-- outFilterMismatchNoverLmax 0.1” to increase the maximum ratio of mismatches to mapped length to 10% (Dobin et al., 2013). The number of reads aligning to genes was counted with featureCounts 2.0.2 tool from the Subread package (Liao et al., 2014). Only reads mapping at least partially inside exons were admitted and aggregated per gene. Reads overlapping multiple genes or aligning to multiple regions were excluded. Differentially expressed genes were identified using DESeq2 version 1.30.0 (Love et al., 2014). Only genes with a minimum fold change of +-2 (log2 = +-1), a maximum Benjamini-Hochberg corrected p-value of 0.05, and a minimum combined mean of 5 reads were deemed to be significantly differentially expressed.

### Cell transfection with GapmeRs

For knockdown assays, in three different biological replicates of human umbilical vein endothelial cells were seeded and allowed to reach 40-50% confluency on the day of transfection. Subsequently, cells were transfected with 50 nmol/L GapmeR targeting PCAT14 or a negative control GapmeR (Qiagen, Paris, France) using Lipofectamine RNAiMAX (Invitrogen, Thermo Fischer Scientific, Dreieich, Germany) according to the manufacturer’s protocol. After 4 hours, the transfection mix was replaced by endothelial growth media and the cells were incubated for additional 48 hours.

### RT-qPCR

Total RNA was extracted using RNeasy kit (QIAGEN, Hilden, Germany) according to manufacturer instructions. Equal amounts of total RNA (1 μg) were reverse transcribed using Superscript III (Invitrogen) according to instructions. Gene expression levels were detected using SYBR Green (Absolute QPCR SYBR Green Mix; Thermo Fisher Scientifc). The relative expression levels of the different genes studied was calculated using the formula 2− □ *C*t (□ *C*t = *C*t (gene) – *C*t (housekeeping gene)) with the 18S RNA as a reference. The primer sequences used were as follows: 18s : forward 5’
s-CTTTGGTCGCTCGCTCCTC-3′, reverse 5′-CTGACCGGGTTGGTTTTGAT-3′ and PCAT14: forward 5′-GGCGCAGGCCACTCCATCTGGTG-3′, reverse 5′-CCTCAATGACCACACTGTAGAG-3′.

### Single molecule RNA imaging (smRNA FISH)

43 oligonucleotides labeled with Quasar 670 targeting human PCAT14 exon sequence were designed with the Stellaris RNA FISH probe designer and produced by LGC Biosearch Technologies. Human umbilical vein endothelial cells were seeded on glass coverslips coated with fibronectin (25 μg/mL) and gelatin (0.5% w/v, Sigma, Darmstadt, Germany). SmRNA FISH was performed as previously described (Dumbovic et al., 2018; Dumbović et al., 2021). Samples were left to curate at 4°C overnight before proceeding to image acquisition.

### Microscopy and image analysis

Image acquisition was performed using a Nikon ECLIPSE Ti2 widefield microscope with an Nikon Plan Apo λ 100x/1.45-numerical aperture oil objective lens and a Nikon DS-Qi2 camera. Z-stacks of 180 nm step size were acquired. Maximum intensity projections were created using ImageJ/Fiji. FISH spots were quantified manually using ImageJ/Fiji, wherein the brightness and contrast of each channel were adjusted.

### Cell proliferation/confluency

Human umbilical vein endothelial cells were seeded onto 24 well plates (Sarstedt, Germany) at a density of 15,000 cells/well. 16 images/well at magnification 10x were taken every 4 hours using an IncuCyte S3 live-cell analysis system (Essen BioScience, United Kingdom). Cell confluence was measured for up to 4 days. Image processing and analysis were performed using the following parameters: segmentation adjustment 1.8; clean up: hole fill 0 μm^2^; adjust size (pixels): 0; filter: minimum 350 μm^2^ area, no maximum, Eccentricity: no minimum, no maximum

### Wound scratch assay

Human umbilical vein endothelial cells were seeded onto 8 well μ-slides (Ibidi) at a density of 40.000 cells/ well. The cell monolayer was scraped in a straight line with a p200 pipet tip 24 hours later and the media was immediately replaced with fresh endothelial growth media. Pictures were taken at 0 and 6 after the wound formation using Axio Observer A1 microscope (Zeiss Oberkochen, Germany) with an AxioCam MRm camera (Zeiss). In order to maintain the same area during the image acquisition marks were created as reference points close to the scratch.

### Spheroid sprouting assay

Spheroids containing 500 cells were generated as described (Korff et al., 2006), and cultured in EBM supplemented with 2.5% FCS and VEGF (30 ng/mL). Phase contrast images were taken after 24 hours and sprouting was quantified by measuring the cumulative length of the sprouts originating from an individual spheroid. Imaging was performed using an Axio Observer A1 microscope (Zeiss Oberkochen, Germany) with an AxioCam MRm camera (Zeiss). At least five spheroids per group were analyzed for each cell batch.

### Monocyte adhesion

Confluent human umbilical vein endothelial cells (cultured in μ-Slide 8 Well, ibidi GmbH, Germany) were subjected to transfection or stimulated with IL-1β (10 ng/ml, 4 hours). Thereafter, THP-1 cells (100.000) were added and left undisturbed for 30 minutes as previously described (Bibli et al., 2019). Non-adherent cells were removed by washing three times with 1x phosphate buffer saline supplemented with 1mmol/L Ca^2+^ and 1mmol/L Mg^2+^ and adherent cells were imaged with Axio Observer A1 microscope (Zeiss Oberkochen, Germany) with an AxioCam MRm camera (Zeiss).

### Data analysis and bioinformatics

To generate a PCAT14-mRNA prediction network, LNCipedia version 5.2, ENCORI for RNA interactomes and NPInter v4.0 (Teng et al., 2020), were used to retrieve experimentally verified mRNAs that could interact with PCAT14. The mRNAs included in the RNAseq data set from the native human endothelial cells were used in the interaction network. PCAT14-miRNA-mRNA prediction network was designed using Cytoscape software.

All graphs were generated using GraphPad Prism 9. Volcano plots were made with EnhancedVolcano package from Bioconductor version 3.15. Hierarchical clustering heatmaps were constructed using RColorBrewer and pheatmap libraries on R software (Clustering method = “average”, clustering distance in rows and columns = “euclidean” or “correlation”).

### Statistics

Data are expressed as mean ± SEM. Statistical evaluation was performed using Student’s t test for unpaired data or paired data. Statistical tests are described in the figure legend for each experiment. Values of P<0.05 were considered statistically significant.

## Results

### Endothelial aging impacts on lncRNA expression profiles

It is well established that lncRNA expression is highly cell type specific (Cabili et al. 2011; Derrien et al. 2012; Molyneaux et al. 2015). To study the expression profiles of lncRNAs in the human native endothelium and analyze the impact of aging on the expression of lncRNAs, we performed RNA-sequencing (RNAseq) on native young (average age 20 years) and aged (average age 80 years) arterial endothelial cells. We 4463 lncRNAs (Table 1). Hierarchical clustering defined differential expression patterns of both mRNAs and lncRNAs during aging and revealed an age-dependent alteration in the transcriptome of the human endothelium (**Figure 1A, 1C**). Transcripts encoding proteins related to endothelial homeostasis and integrity (i.e., PLXDC2, NOS1, ANGPT1, FGFR2 and CDH23) as well as metabolic related transcripts (i.e., LARGE1, CYP7B1 and CYP1B1) were downregulated during aging, whilst genes associated with disease, apoptosis and inflammation were upregulated (i.e., BBC3, CXCL6, CDKN2A and BNC1) (**Figure 1B**). We observed a significant transcriptional reprogramming of lncRNAs, with 601 lncRNAs being downregulated (≤-1 Log_2_ fold change, p<0.05) and 197 lncRNAs being upregulated (≥1 Log_2_ fold change, p<0.05) in aged human endothelium (**Figure 1D**). To determine whether the differentially expressed (DE) lncRNAs originate from specific chromosomes and chromosome regions, which would therefore reflect epigenetic landscape changes known to occur in upon aging (Ding et al., 2020), we analyzed the distribution of affected lncRNAs in the genome. The differentially expressed lncRNAs were distributed among the chromosomes with of the majority of them positioned on chr1, chr2, chr6, chr8, chr9 and chr11 (**Figure 1E**). Overall, our analysis shows that expression of lncRNAs is significantly affected by aging in primary endothelial cells, with many of them being involved in regulation of endothelial cell function.

**Figure 1.**
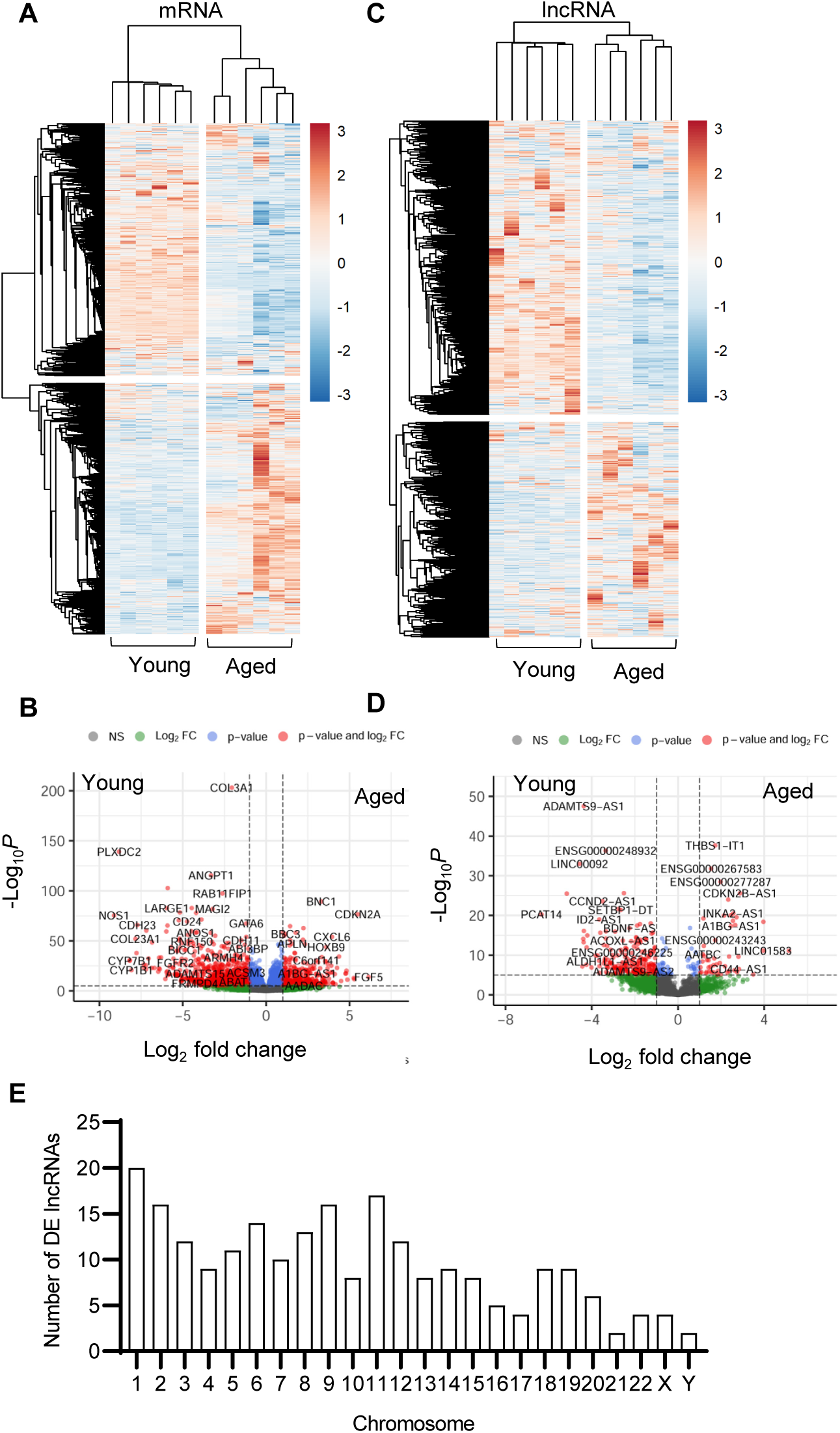
Effect of aging on endothelial cell transcriptome. Bulk RNA-sequencing analysis was perfomed in human native young and aged arterial endothelial cells (n=6 per group, each sample is a pool of 5 different isolates). (A,C) Hierarchical clustering heatmap showing differentially expressed mRNAs (A) and lncRNAs (C). (B,D) Representative volvano plots showing differentially expressed mRNAs (B) and lncRNAs (D) (total mRNA variables = 15379, total lncRNA variables = 4463). (E) Chromosome distribution of differentially expressed (DE) lncRNAs detected as in panels C and D.

### PCAT14: the major endothelial age-dependent downregulated lncRNA

lncRNAs whose expression was most affected in the aged human endothelium compared to endothelial cells isolated from young arteries were PCAT14, LINC00092 and ADAMTS9-AS1 (**Figure 2A**). Given that the lncRNA PCAT14 was among the most highly abundant expressed lncRNAs in the native human endothelium shown to be reduced in the aged endothelial cells, we focused our further studies on PCAT14. A detailed analysis of PCAT14 locus revealed interesting and distinguishing features: i) PCAT14 is not in a vicinity of any protein coding gene, the nearest one being IGLL1 (Immunoglobulin lambda-like polypeptide 1) located approximately 25 kb downstream of the PCAT14 locus; and ii) PCAT14 locus overlaps with a human specific endogenous retrovirus H element (HERVH), therefore making it a primate-specific, transposon-derived lncRNA (**Figure 2B**). By quantitative RT-PCR (RT-qPCR) we validated its relatively high expression in young endothelial cells and significant reduction in expression in aged endothelial cells (**Figure 2C**). We used single molecule RNA imaging (smRNA FISH) targeting PCAT14 exons to analyze the distribution of PCAT14 lncRNA in human native endothelial cells isolated from individuals of different age. Our analysis shows that PCAT14 is almost exclusively a nuclear lncRNA, with most clear signal forming bright punctuate foci (**Figure 2D**). Interestingly, these foci are most frequently localized at the nuclear periphery or lamina, which are regions known to consist of heterochromatin.

**Figure 2.**
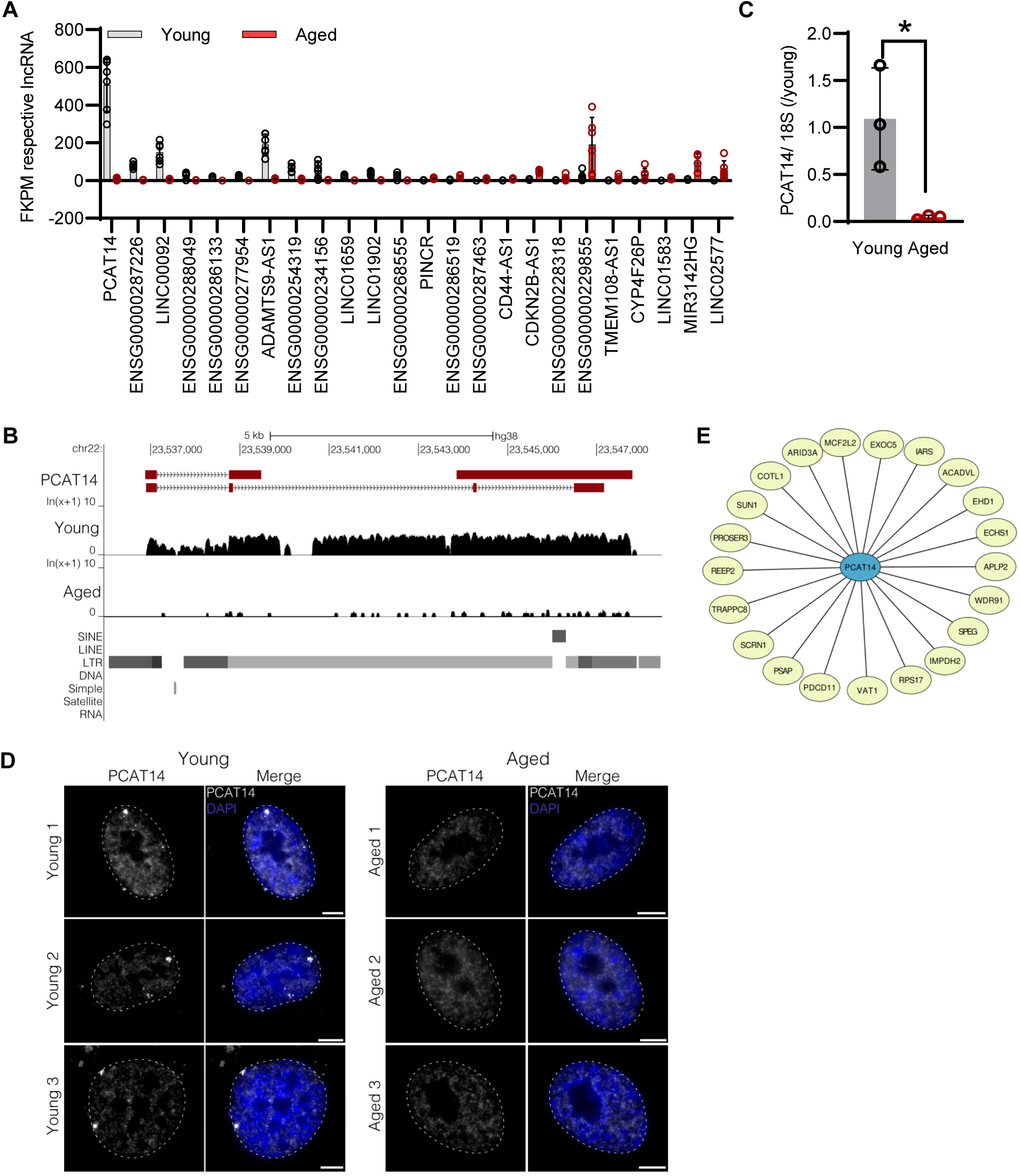
Impact of aging on endothelial PCAT14 expression, localization and mRNA network. (A) Expression levels (FPKM) of lncRNA transcripts in samples as in Figure 1. (B) UCSC Genome Browser showing the PCAT14 locus (hg38) and the poly(A)+ RNA-seq tracks from young and aged human native endothelial cells. (C) Relative lncRNA levels of PCAT14 in young and aged native isolated arterial endothelial cells (n=3 per group,*P<0.05, Student’s t-test). (D) Maximum intensity projections of representative images of PCAT14 smRNA FISH on young and aged human native endothelial cells. Exon, gray; nucleus, blue, outlined with a dashed circle. bar = 5 μm. RNA FISH was performed on 7 young and 4 aged samples. (E) Prediction network of PCAT14 with mRNAs (yellow nodes).

Next, we constructed a co-expression interaction network to link PCAT14 with all the potential mRNAs appearing in our bulk RNA-sequencing dataset. Such a network could provide an essential framework for mapping sets of mRNA and lncRNAs that could act in a coordinated manner. Interestingly, PCAT14 had a potential regulatory relationship with several protein-coding genes related to endothelial metabolism i.e. ACADVL, (Schoors et al., 2015), development i.e. ARID3a (An et al., 2010) and junctional integrity i.e. EHD1 (Deracinois et al., 2012), indicating that PCAT14 potentially plays a role in the maintenance of the vascular fitness (**Figure 2E**).

### PCAT14 regulates key endothelial cell functions

To address the role of PCAT14 in endothelial cells, we used human umbilical vein endothelial cells (HUVEC) transfected with a GapmeR against PCAT14 (PCAT^KD^) or a control GapmeR (CTL). By RT-qPCR we show that PCAT14 GapmeRs successfully downregulated its expression by approximately 60% (**Figure 3A**). Downregulation of PCAT14 did not affect the endothelial proliferative capacity (**Figure 3B**). However, diminished PCAT14 expression reduced the endothelial cell migratory ability approximately by 30% (**Figure 3C**) and reduced the sprouting capacity by approximately 35% (**Figure 3D**). These data indicate that PCAT14 preserves endothelial regenerative and angiogenic properties. Finally, we investigated the pro-inflammatory phenotypes of the PCAT^KD^ endothelial cells through an endothelial cell-monocyte adhesion assay. Human monocytes (THP-1) exhibited increased capacity to adhere to endothelial cells lacking PCAT14, to a similar extent as endothelial cells stimulated prior with interleukin 1 beta (Il-1β) for 4 hours, indicating that PCAT14 impacts on endothelial cell activation (**Figure 3E**).

**Figure 3.**
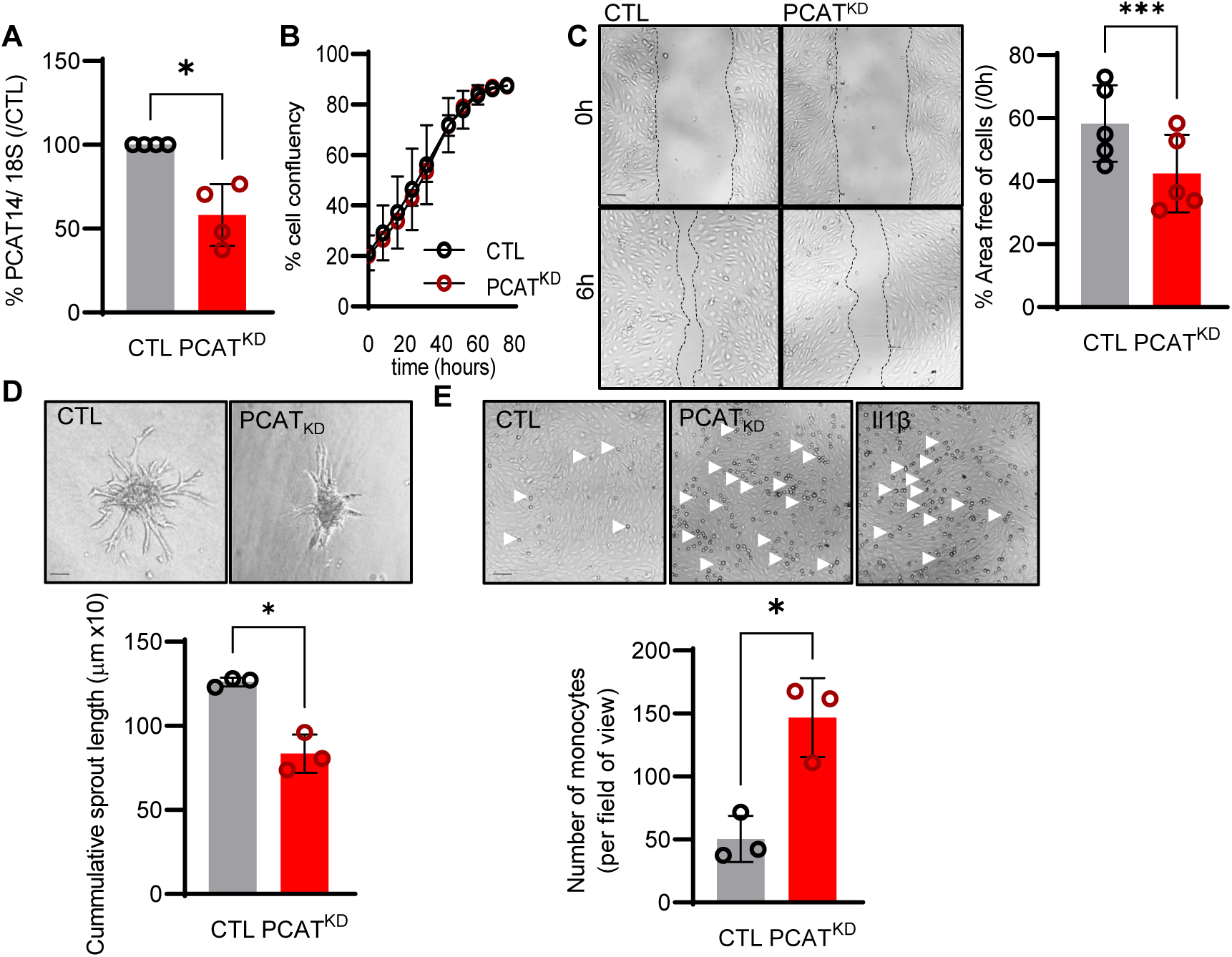
Effect of PCAT14 deletion on endothelial angiogenic responses and activation. Human umbilical vein endothelial cells (HUVEC, Passage 3) were transfected with GapmeR (50nmol/L, 72h) targeting PCAT14 (PCAT^KD^) or a respective control (CTL). (A) Relative PCAT14 expression normalized to CTL. (B) Percentage of endothelial cell confluence for up to 48 hours. (B) Representative brightfield images and quantification following a wound scratch assay at 0 and 6 hours. (D) Representative brightfield images and quantification of endothelial cell sprouts on a 3D spheroid assay. (E) Representative brightfield images and quantification of THP-1 monocytes adhered to endothelial cells. As positive control endothelial cells treated with Interleukin 1β (Il-1β, 10ng/mL, 4 hours). (n=3-6 biological replicates per group, experiments performed at least 2 times. bar = 500μm, *P<0.05, **P<0.01, Student’s t-test paired).

## Discussion

Our data highlight that aging alters the endothelial cell lncRNA expression profiles. We identified 798 lncRNAs being significantly altered in the aged human native endothelium. Focusing on one of the most endothelial abundant and simultaneously age sensitive lncRNA, PCAT14, it was possible to link its expression with the preservation of endothelial cell migration, sprouting as well as anti-inflammatory potential.

Advanced aging is an independent risk factor related with the development of endothelial dysfunction even in the absence of a clinical disease. With advanced age the endothelium is characterized by increased inflammation, oxidative stress and senescence (Campisi and Di d’Adda Fagagna, 2007). Endothelial cell senescence is coupled with genomic instability, telomere attrition and DNA damage which are able to trigger the p53/p21 pathway and eventually lead to cell cycle arrest (van Deursen, 2014; Donato et al., 2015). Notably, the elimination of senescent cells (senolysis) was recently reported to improve normal and pathological changes associated with aging in mice (Baker et al., 2016; Xu et al., 2018) and the recent development of endothelial cell-specific senolytic targeting seno-antigens could be a promising strategy for new senolytic therapies to diminish pathological phenotypes associated with aging (Suda et al., 2021).

Although extensive work has been done to identify and characterize lncRNAs in multiple diseases (He et al., 2018), including cellular senescence (Bink et al., 2019), only a few have been examined for their potential actions in the endothelial aging and age-associated vascular diseases. Among those linked to vascular aging, MALAT1, H19 and GFADS5 have been shown to impact endothelial function and induce aging by modulating the expression of the cell cycle regulators (Michalik et al., 2014), or regulate the endothelial epigenome and its angiogenic responses (Hofmann et al., 2019) (Jiang and Ning, 2020). The lncRNA Mirial was recently shown to be downregulated in the aged murine endothelium and impact on endothelial metabolic and cellular functions (Kohnle et al., 2021). The lncRNA ANRIL seems to be increased in advanced endothelial aging both in murine models (Tsai et al., 2010) and patients with coronary artery disease, atherosclerosis and type 2 diabetes. Its actions have been attributed on its ability to bind either on p300 and affect VEGF expression or on polycomb repressive complex-2 (PRC2) complex (Thomas et al., 2017) and disturb its binding to the p15^INK4B^ locus (Kotake et al., 2011). The lncRNA HOTAIR is considered to affect also the PRC2 and the lysine-specific histone demethylase-1 complexes and affect the levels of H3K27me3 and H3K4me2 (Tsai et al., 2010), inducing atherosclerotic events during aging (Pierce and Feinberg, 2020). The lncRNAs Meg3 and lincRNA-p21 affect p53 expression and are linked to vascular aging (Huarte et al., 2010; Shihabudeen Haider Ali et al., 2019). The current literature links the aformentioned lncRNAs with endothelial cell function and development of age related characteristics or altered endothelial cell signaling responsible for the development of age related phenotypes. In our datasets we were able to validate that in the native human aged endothelium the lncRNA H19 was significantly downregulated and ANRIL was significantly upregulated. MALAT1, Meg3, GFADS5 and HOTAIR were indeed expressed in the human native endothelium, however, their expression profiles seem to be age independent. This might be attributed to the fact that studies investigating these lncRNAs rely on the *in vitro* induced endothelial replicative senescence is mostly related to the telomere attrition and it can only partially mimic the *in vivo* complex phenomena taking place in the progression of cellular aging. An alternative explanation would be that the current literature investigates the expression profiles of the aformentioned lncRNAs in the presence of additional co-morbidities i.e. atherosclerosis, diabetes or carotid artery diseases, which sometimes precede endothelial senescence.

lncRNAs identified as being the most downregulated in the human native aged endothelium have been described in the context of cancer. In particular ADAMTS9-AS1, is highly expressed in hepatocellular carcinoma cells and it exacerbates cell proliferation, migration and invasion through the PI3K/AKT/mTOR pathways (Zhang et al., 2020). LINC00092 affects the glycolysis levels and modifies the Warburg effects in ovarian cancer progression and metastasis (Zhao et al., 2017). Given that endothelial cells heavily rely on glycolysis, further studies investigating LINKC000092 in the context of endothelial rejuvenation, would be promising. The lncRNA CCND2-AS1 has been identified in papillary thyroid carcinoma, breast cancer and cervical cancer and promotes proliferation, migration and invasion (Chen et al., 2018). In contrast to the most down-regulated lncRNAs which are linked to cell proliferation, migration and invasion the most upregulated lncRNAs were linked to senescence. In particular, THBS1-IT1, highly expressed in the centenarian population, has been linked to decrease of p16, p21 and the activity of the senescent related beta-galactosidase (Jiang et al., 2021).

We focused our further studies on PCAT14 due to its high expression in native young endothelium which was not detected in the aged endothelium. PCAT14 is mostly expressed in the prostate and the testis and less in the thyroid, lung and ovary. At the same time, PCAT14 has been characterized as a novel prostate cancer and lineage-specific lncRNA (Shukla et al., 2016; Yan et al., 2021). In the native human endothelium, PCAT14 was one of the most abundant age sensitive lncRNAs. An interesting observation is that PCAT14 originates from a HERVH, a primate-specific endogenous retrovirus. HERVH has been of great interest due to its high expression in both human embryonic stem cells (hESCs) and induced pluripotent stem cells (iPSCs) (Loewer et al., 2010; Kelley and Rinn, 2012; Santoni et al., 2012; Fort et al., 2014; Wang et al., 2014; Zhang et al., 2019). The relationship between HERVH and pluripotency has been well established. Knockdown of HERVH family results in the loss of hESC identity and reduced reprogramming efficiency to pluripotency (Lu et al., 2014; Ohnuki et al., 2014; Wang et al., 2014). Knocking down individual HERVH-derived lncRNAs, such as lincRNA-RoR, or deleting individual HERVH loci acting as topologically associated domain (TAD) boundaries leads to similar phenotypes (Loewer et al., 2010; Wang et al., 2014; Zhang et al., 2019). Overall, these studies support the notion that HERVH products regulate the cellular homeostasis of pluripotent stem cells with several hypotheses put forward for their regulatory role, including acting as enhancers, lncRNAs, and TAD boundaries.

In our smRNA FISH analysis on native endothelial cells from young and aged individuals, we observed PCAT14 as almost exclusively nuclear lncRNA in native young endothelium. The distribution of PCAT14 points towards it functioning *in cis*, while the extensive silencing of PCAT14 locus upon aging suggests epigenetic reprograming of the locus upon aging. Important to note, although we detect PCAT14 expressed in young endothelium, it is normally silenced in differentiated tissues. In line with demonstrated functions of HERVH-derived lncRNAs in stemness, the high expression of PCAT14 in young endothelial cells may point towards its potential role in endothelial regenerative capacity. Further studies will be needed to determine the exact molecular mechanism of PCAT14 lncRNA and the regulation of the locus. mRNA-PCAT14 prediction network, predicted PCAT14 interactions with mRNAs which are linked to endothelial disease and vascular aging. The mRNA SUN1 encodes for the inner nuclear envelope protein SUN1, a component of the linker of nucleoskeleton and cytoskeleton complex. Sun1 modulates endothelial peripheral cell-cell junctions from the nucleus and impacts vascular development and barrier function (Bouzid et al., 2019; Buglak et al., 2021). Another interesting potential interactor is the very long-chain specific acyl-CoA dehydrogenase (ACADVL), which holds a role in the fatty acid oxidation metabolic pathway and whose silencing of in the endothelium impairs vessel sprouting (Schoors et al., 2015). Perhaps the most interesting predicted PCAT14-mRNA interaction is with AT-rich interaction domain 3A (ARID3A). ARID3A levels are increased in patients with pulmonary artery hypertension (PAH) (Reyes-Palomares et al., 2020). The protein is related to endothelial activation and has been shown to regulate p53 functions (Lestari et al., 2012). Given such interactions, we investigated whether PCAT14 would have potential impact on endothelial cell migration, sprouting and inflammatory responses. Indeed, by knocking down PCAT14 in young endothelial cells during proliferation we observed diminished migratory capacity, reduced sprouting as well as increased activation.

Taken together, our data highlight that aging can strongly affect lncRNA expression profiles, which may potentially affect endothelial fitness. Downregulation of one such age-sensitive lncRNA PCAT14 affects major endothelial functions. Therefore, going forward, it will be important to investigate the individual and/or combined roles of lncRNA expression alterations and mechanisms thereof on endothelial cell functions.

## Data availability statement

The authors declare that the data supporting the findings of this study are available within the paper.

## Ethics statement

The use of human material in this study conforms to the principles outlined in the Declaration of Helsinki and the isolation of endothelial cells was approved in written form by the ethics committee of the Goethe-University.

## Author contributions

Study conceptualization and design: S.-I.B., G.D.; experiment design, performance, and data analysis: S.-I.B., G.D., M-K.D. and R.C.E.; computational analyses: S.G., M.L. and S.T.; funding: S.-I.B. and G.D., writing the paper: S.-I.B., G.D. and M.-K.D.; all authors have read and approved the manuscript before submission.

## Acknowledgements

The authors are indebted to Cindy Fabiene Hoeper for expert technical assistance.

